# Geometric morphometric analysis of facial sexual dimorphism in a contemporary sample : An application to sex prediction of ancient human remains

**DOI:** 10.1101/2025.06.17.660135

**Authors:** Juliette Legros, Pauline Borde, Frédéric Savall, Fabrice Dédouit, Eric Crubézy, Norbert Telmon

## Abstract

Focusing on craniofacial bones, this study investigates morphological variation related to sexual dimorphism in order to deepen our understanding of human biological diversity and to provide new data from a contemporary reference sample. Accordingly, the research was guided by three objectives (i) identify the facial regions exhibiting the greatest sexual dimorphism using landmark-based geometric morphometric method; (ii) evaluate the reliability of discriminant models based on these dimorphic regions; and (iii) conduct exploratory analyses to assign sex classification probabilities to ancient subjects using the discriminant models derived from a contemporary reference sample. The reference sample comprised 44 skulls from subjects of known sex who died in 2024. The ancient sample included 4 skulls recovered from the Grotte de La Medecine (France), attributed to the chalcolithic period. Fourteen facial landmarks were digitized using 3DSlicer. Generalized Procrustes Analysis was performed to extract shape variables and standardized coordinates for statistical analysis. Thin -Plate Spline transformations quantified and visualized deformation amplitudes between the female and male shapes. Landmarks in the orbital showed the highest deformation amplitudes. Goodall F-test comparing male and female shapes across three facial regions revealed significant sexual dimorphism only in the orbital region and the global facial shape. Discriminant analysis demonstrated that the orbital region provided the highest classification accuracy (88.6%) compared to the global facial region (72.7%). The discriminant models yielded high probabilities of male and female sex classification for the ancient subjects. Using a reproducible method, comparable levels of accuracy to those reported in the scientific literature were achieved despite a relatively modest sample size. These findings further confirm the significant role of the orbital region in human craniofacial sexual dimorphism.

## Introduction

The establishment of the biological profile is a crucial step in the analysis of human remains, as it provides information on an individual’s age, sex, ancestry, stature, and health condition (Ubelaker 1989; Bruzek & Murail 2006). It has applications in numerous fields (White et al. 2011). In bioarchaeology, sex estimation is essential for studying paleodemography and enables the interpretation of various cultural aspects, such as the division of labor or funerary behavior (Lozano et al. 2021; Rivolla 2023). In forensic science, establishing the biological profile is necessary for individual identification (Garvin 2012). Among these parameters, sex estimation holds particular importance as it conditions the interpretation of the other characteristics of the biological profile (Delabarde and Ludes 2014; Ubelaker & DeGaglia 2017).

Sex estimation requires a prior understanding of anatomical variations between male and female subjects, termed sexual dimorphism. The pelvic bone is widely regarded as the most reliable element of the human skeleton for sex determination, with reported accuracies exceeding 90% (Phenice 1969; Sutherland & Suchey 1991; Ubelaker & Volk 2002; Murail et al. 2005). However, in many forensic or archaeological contexts, the pelvis may be absent, poorly preserved, or too fragmented, requiring the use of alternative skeletal elements for sex estimation (Bruzek 1992). Numerous studies have highlighted the diagnostic potential of the skull (White & Folkens 2005; Iscan & Steyn 2013; Ogawa et al. 2013; Marinescu et al. 2014). Cranial methods typically rely on a combination of morphoscopic traits scored on a scale from 0 to 5, corresponding to anatomical expressions categorized as “undetermined,” “female,” “probable female,” “probable male,” “male,” and “ambiguous” (Buikstra 1994). Alternatively, morphometric approaches based on sets of craniometric measurements are frequently employed (Howells 1973; 1989; 1995). However, both approaches are subject to limitations: the inherent subjectivity involved in scoring trait expressions and a high degree of dependence on the observer’s experience and training (Langley and Jantz 2020; Ubelaker & DeGaglia 2020).

Geometric morphometrics has emerged as an innovative approach in biological anthropology. Although size is a major factor in craniofacial sexual dimorphism and has been thoroughly characterized through morphometric techniques, anatomical differences between males and females also arise from shape-specific traits. The advancement of geometric morphometric methods in recent years has significantly enhanced the ability to explore these shape-based variations independently of size (Bookstein 1996; Slice 2005). This methodology is based on the analysis of the three-dimensional coordinates of homologous anatomical landmarks, enabling the quantitative study of shape independently of orientation, nor position of the structures. Shape variations are visualized using Thin-plate spline (TPS) deformation grids. These transformations interpolate shape changes based on the spatial configuration of landmark coordinates. These grids illustrate localized expansions or contractions across individuals, in our case, between the average male and female shapes. Moreover, the amplitude of deformation corresponds to the bending energy required to transform the reference shape into the target shape, providing a quantitative measure of morphological change (Bookstein 1990; Webster and Sheets 2010). Predictive approaches based on models such as linear discriminant analysis (LDA), logistic regression, and support vector machines (SVM) are commonly used to develop functions for sex estimation (Santos et al. 2014; Ogawa et al. 2013; Del Bove et al. 2020). These methods classify shapes by estimating the probability of an individual being male or female. Although various methods are available, the present study relied exclusively on LDA.

The integration of geometric morphometrics with imaging techniques such as postmortem computed tomography (PMCT) offers a non-destructive approach for the analysis of skeletal remains. This method is particularly valuable in archaeological contexts, where conservation constraints often restrict invasive interventions (Dedouit et al. 2014; Uldin 2017; Bédécarrats 2023). PMCT was employed in the study of human remains recovered from the Grotte de La Médecine. This archeological site discovered in 1959 by Jacques Pomié in Verrières (Aveyron, France) was excavated twice in 1960, and yielded numerous skeletal remains, including crania preserved within calcite concretions (Pomié 1967; Soutou 1967). The skulls have been assigned to the Chalcolithic period, ca. 2600–2300 BCE, due to the presence of archaeological material (copper axe) found alongside them (Soutou 1967; Balsan & Costantini 1972). The presence of a calcite matrix around the bones restricted direct examination, thereby justifying the use of non-destructive imaging techniques to virtually access and study the skeletal remains while preserving their physical integrity. Remains recovered in such condition were therefore scanned, virtually cleaned, and reconstructed in 3D using the software 3D Slicer. Among these human remains four skulls were selected to be studied.

The present study focuses on three objectives: (i) apply geometric morphometric methods to quantitatively describe facial sexual dimorphism within a contemporary French sample. Our approach is based on both a “global” and a “local” analysis. In this context, “global” refers to the overall craniofacial shape, encompassing all recorded landmarks, whereas “local” pertains to more restricted anatomical regions defined by a limited subset of landmarks corresponding to specific facial areas. (ii) Develop and evaluate discriminant functions based on facial regions exhibiting significant sexual dimorphism. (iii) Assess whether the regions exhibiting significant sexual dimorphism could be used to predict the sex of ancient subjects from the *Grotte de La Médecine,* using discriminant functions developed on facial shapes belonging to the reference sample.

## Materials and Methods

### Sample

The reference sample was created from post-mortem cranial scans from 44 individuals, 22 females (mean age = 47.9 years, maximum age = 86 years, minimum age = 20 years, and standard deviation = 17.6 years) and 22 males (mean age = 43.5 years, maximum age = 76 years, minimum age = 20 years, and standard deviation = 16 years) (Fig.1). The reference sample was built using anonymized data. Each individual was assigned a unique identifier prior to post-mortem computed tomography (PMCT), which was then linked to the medico-legal database of Toulouse. This database provides verified information on the age and sex of the subjects, as well as relevant details regarding their health status. Based on these data, subjects who were immature or exhibited trauma or pathological conditions likely to alter facial morphology were excluded from the study.

**Fig 1.**
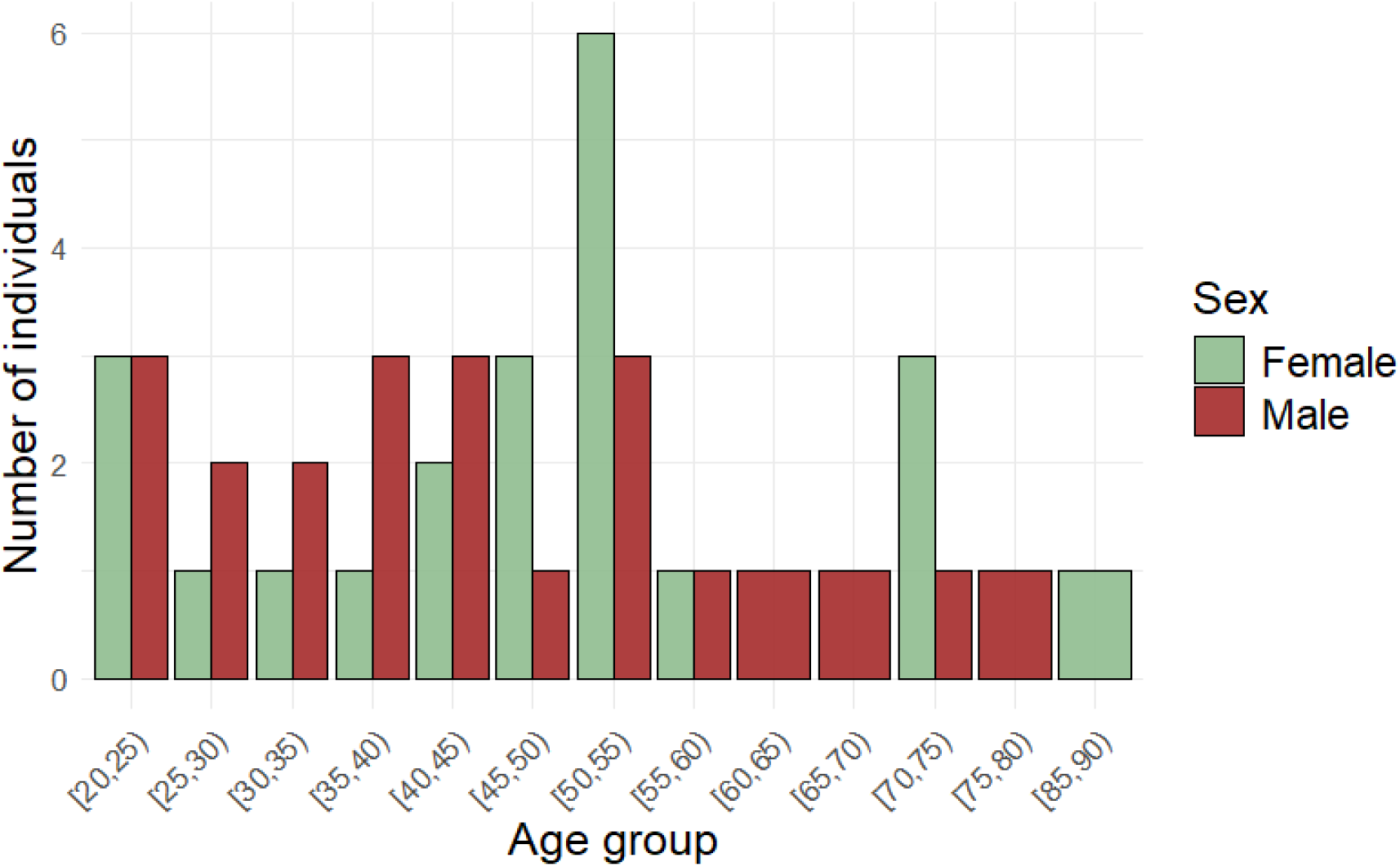
Histogram showing the age distribution of male (M) and female (F) subjects in the contemporary reference sample. Female subjects are represented in green, male subjects in brown.

Regarding the skulls recovered among other skeletal remains in the Grotte de La Médecine (Fig. 2), three criteria had to be met to include a subject in the archaeological sample used in this study.

1. Since sexual dimorphism is difficult to assess in immature subjects (Delabarde & Ludes 2014), a minimum age was estimated to ensure that only adult or subadult skulls were included in the analysis. Age was estimated based on dental eruption, developmental stages and calcification (Ubelaker 1987; AlQahtani 2008). Complete spheno-occipital synchondrosis fusion was also used (Schaefer et al. 2009).
2. The state of preservation was considered. A skull is considered ‘well preserved’ if it shows no taphonomic degradation affecting the anatomical regions involved in this study, or the surrounding regions likely to alter facial shape.
3. No lesions or pathological conditions, considered to be in response to disease or injury (Mann & Murphy 1990), were to be observed.

**Fig 2.**
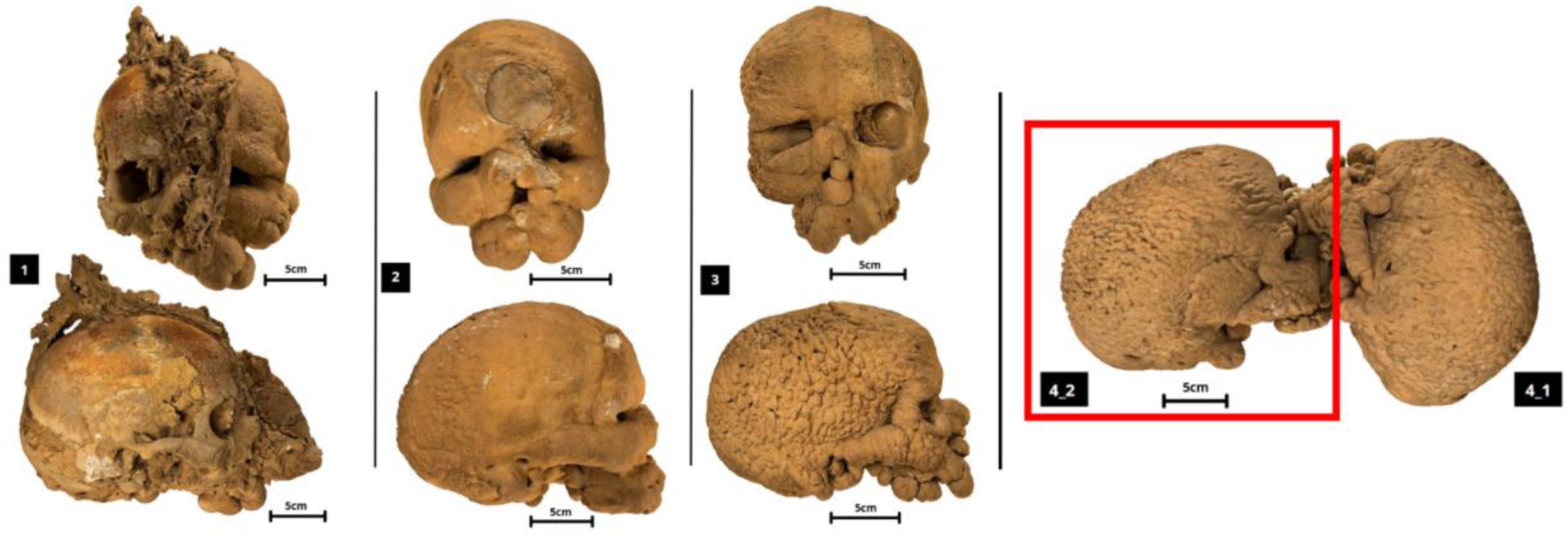
Skulls from the Grotte de La Médecine used in this : subjects 1, 2, 3, 4.2 (‘4.2’ will henceforth be referred to as ‘4’ in this study). Top: anterior view, bottom: right lateral view. Scales corresponding to real 5cm are associated with each skull for each view.

Eight skulls were recovered from the Grotte de la Médecine. Among them, two subjects (labelled 4.1 and 4.2) were encased within the same calcite block. Only subject 4.2 was included in the present study, as subject 4.1 exhibited a maxilla too fragmented to meet the established preservation criteria. For clarity, subject 4.2 is hereafter referred to as subject 4. Of the eight crania initially recovered, only four (subjects 1 to 4) were retained for analysis. In addition to subject 4.1, two were excluded due to their immature status, and two others due to the absence of facial structures, with only partial preservation of the neurocranium.

### Radiological features

For both ancient and contemporary individuals, post-mortem computed tomography (PMCT) scans were performed at the University Hospital Centre of Toulouse, France, at the Institute of Forensic Medicine during 2024 (SIEMENS Somatom, definition AS). The slice thickness was 0.75 mm for contemporary individuals and 0.6 mm for ancient individuals, with a collimation of 0.6 mm, a 512×512 matrix for all and 140 kV for all. Digital Imaging and Communications in Medicine (DICOM) files have been used to store the CT scans.

### Mesh acquisition and landmarks setting

Post-processing was performed with 3D Slicer software (version 5.7.0, 3D Slicer s.d.; Fedorov et al. 2012). 3D models of contemporary individuals have been made using a threshold segmentation method to separate bones from other body tissues. This separation is achieved thanks to their different densities. An additional step was required to process the chalcolithic subjects. They were first virtually cleaned to remove agglomerated calcite from the inside and outside of the skulls (Fig. 3). Because of the overlapping densities of calcite matrix and bone, the threshold separated poorly the two structures. The segmentation was retouched (approximately every 5 slices) to ensure no calcite was included in the bone segment. Once the 3D models were completed, they were used in « .ply » format (polygon file format)

**Fig 3.**
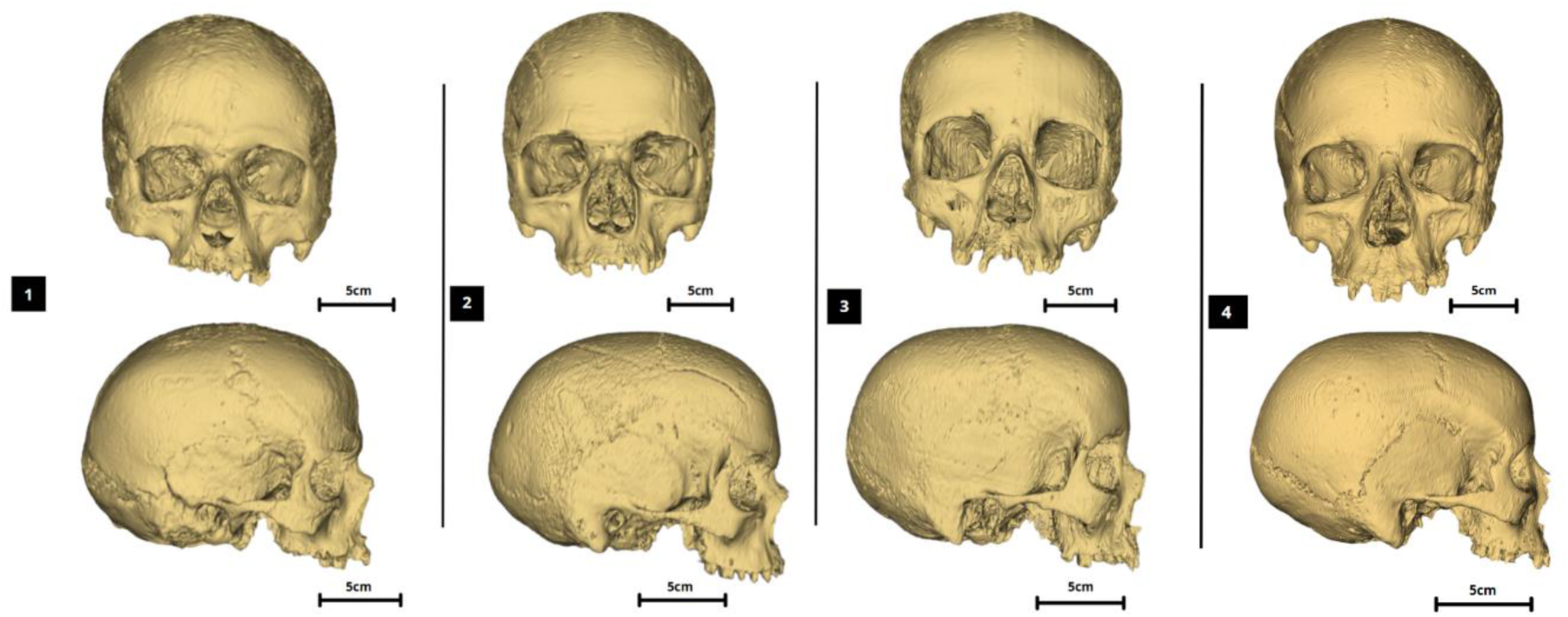
Virtual 3D reconstructions of ancient skulls from the Grotte de La Médecine (subjects 1–4), generated through PMCT imaging and segmentation processing. Top: anterior view, bottom: right lateral view. Scales corresponding to real 5cm are associated with each skull for each view.

Fourteen landmarks of interest (Table 1) in adult facial sexual dimorphism were selected (Fig. 4). 3D models were used to facilitate landmark placement on the skull, but final position was verified using the PMCT slices (in axial/ coronal/ sagittal views) in order to ensure anatomical accuracy, employing the software’s Markups module. The x, y and z landmark coordinates of each subject were recorded in « .json » format.

**Fig 4.**
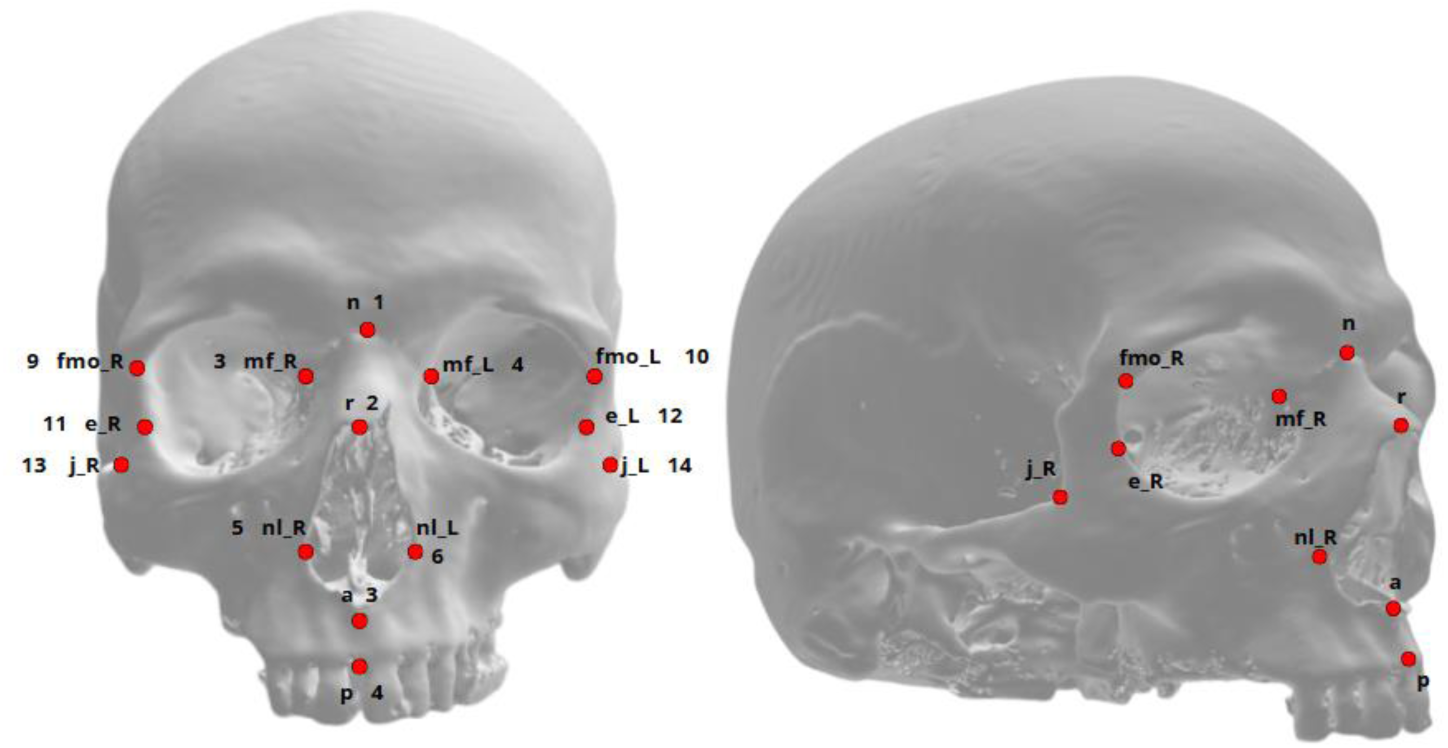
Cranial landmarks chosen for this study placed on a 3D reconstruction of a human skull. Left: anterior view; Right: right antero-lateral view.

**Table 1.**
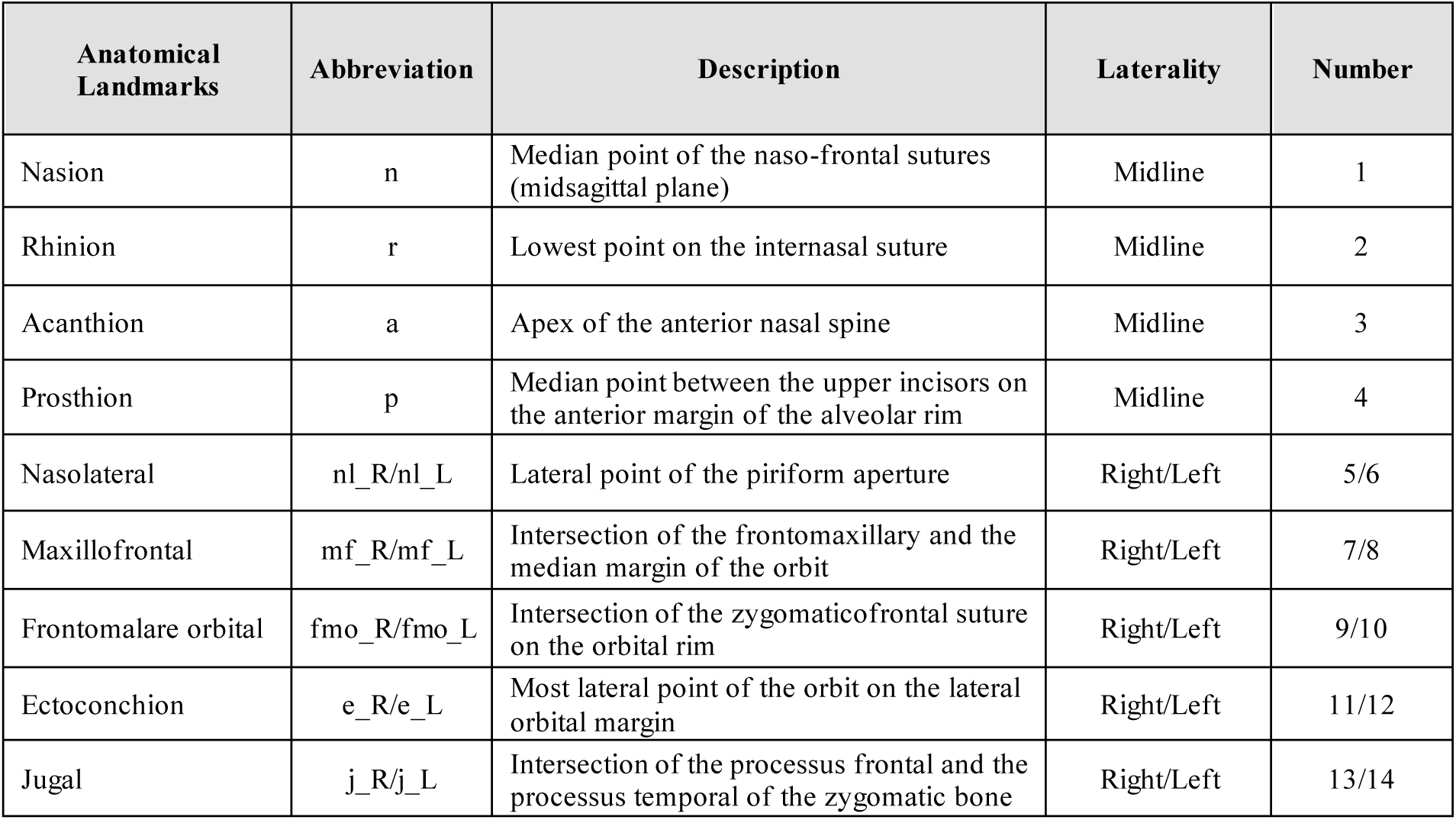
Description of the fourteen facial landmarks.

### Statistical analysis

Statistical analyses were performed using Rstudio software (version 4.4.3 2025; Team R 2015; R Core Team 2021). A p-value of less than 0.05 was considered statistically significant. The ‘ggplot2’ R package was used for 2D plots, and ‘rgl’ for 3D plots.

To assess intra-observer variability, the primary observer repeated the landmark placement one month after the initial session to minimize memory bias. For each subject, Euclidean distances between corresponding landmarks from the two sessions were calculated. For the inter-observer variability assessment, two independent observers placed the same set of landmarks on each subject. Both observers were provided only with a standardized written description of the anatomical landmarks, without access to each other’s work, to ensure independent evaluation. For each landmark, the mean Euclidean distance and percentage error between observers were computed along the three spatial dimensions (x, y, z). In addition, repeatability was quantified by calculating the intraclass correlation coefficient (ICC) for each landmark thanks to the irr R package. High ICC values indicate greater consistency in landmark placement across observers (Table 2).

**Table 2.**
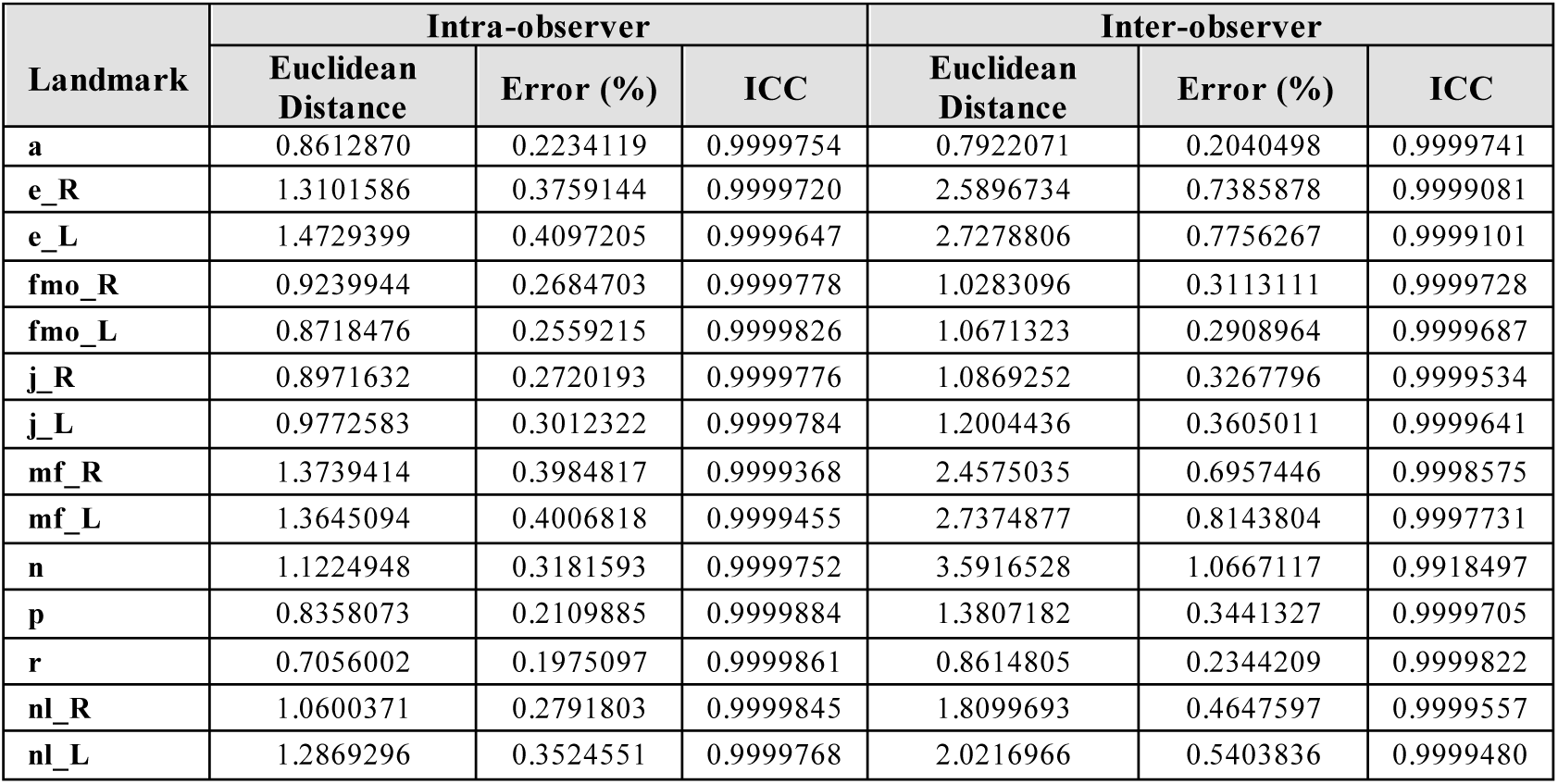
Intra- and inter-observer variability.

### Geometric morphometric analysis

Landmark data standardization was achieved via a Generalized Procrustes Analysis (GPA), over the 48 subjects (44 contemporary and 4 ancients), eliminating the effects of position, and orientation (Slice 2007). As part of this procedure, all specimens were scaled to unit centroid size to ensure that the observed differences were solely related to shape, independent of size.

Principal Component Analysis (PCA) were performed to explore the morphological variation in the human facial shape represented by the 14 landmarks (Fig. 5) and the orbital region (Fig. 6). The PCA reduced the dimensionality of the landmarks Procrustes three-dimensional coordinates (x, y, z) dataset to isolate the components of variation related to shape. Sex variable was used as the statistical explanatory variable for the analysis of shape difference. The sample analyzed is composed of two subgroups: contemporary individuals, of known sex (male and female). The second subgroup corresponds to ancient individuals whose sex remains undetermined; they were subsequently projected into the morphospace defined by the principal components.

**Fig 5.**
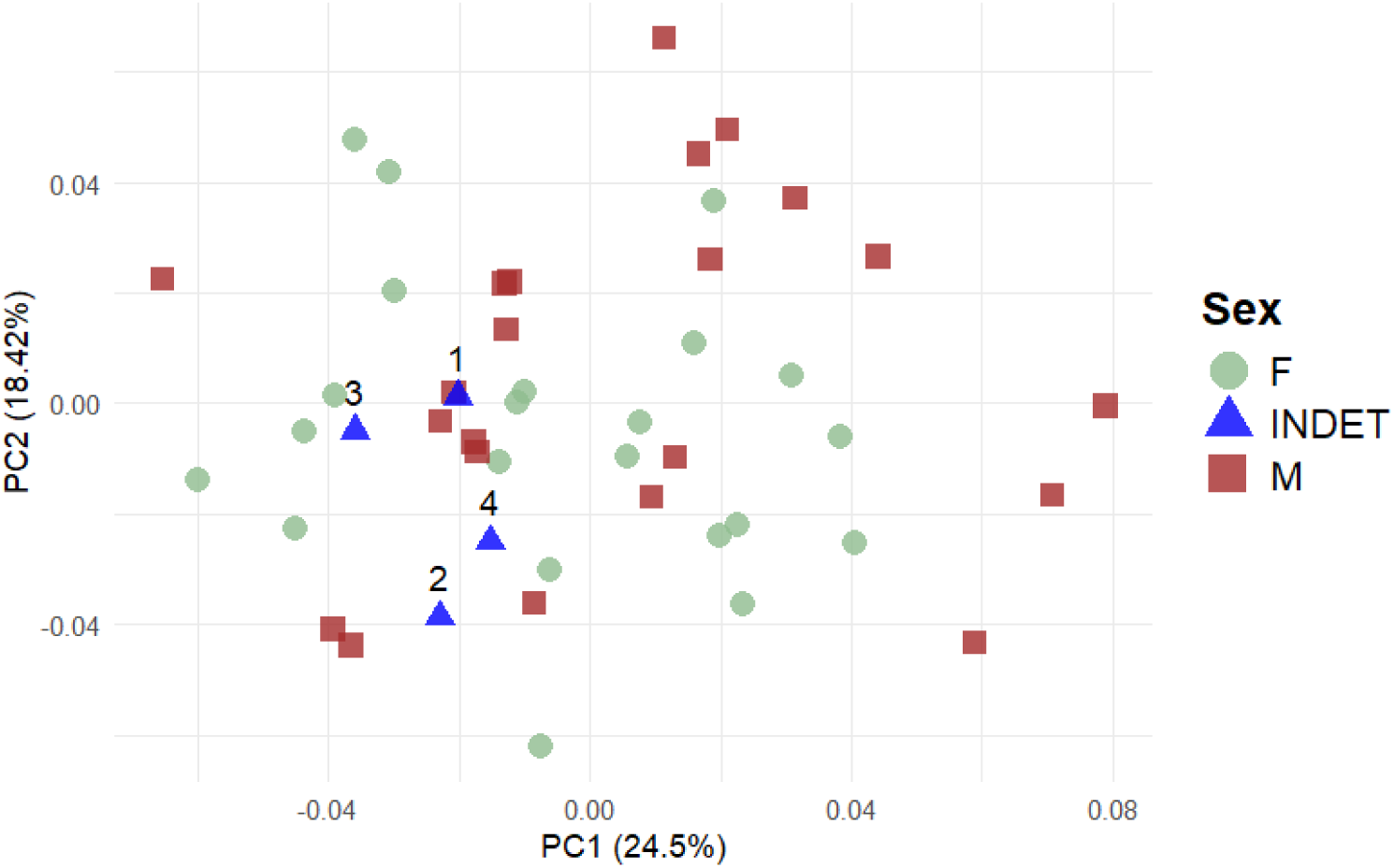
Morphospace is defined by the first two principal components from the PCA of the overall shape of the face traced by the 14 three-dimensional landmarks. Contemporary male skulls are represented by brown square, females by green dots; and ancient skulls of undetermined sex are associated with blue dots. The ID number of each ancient subject is annotated next to the corresponding dot.

**Fig 6.**
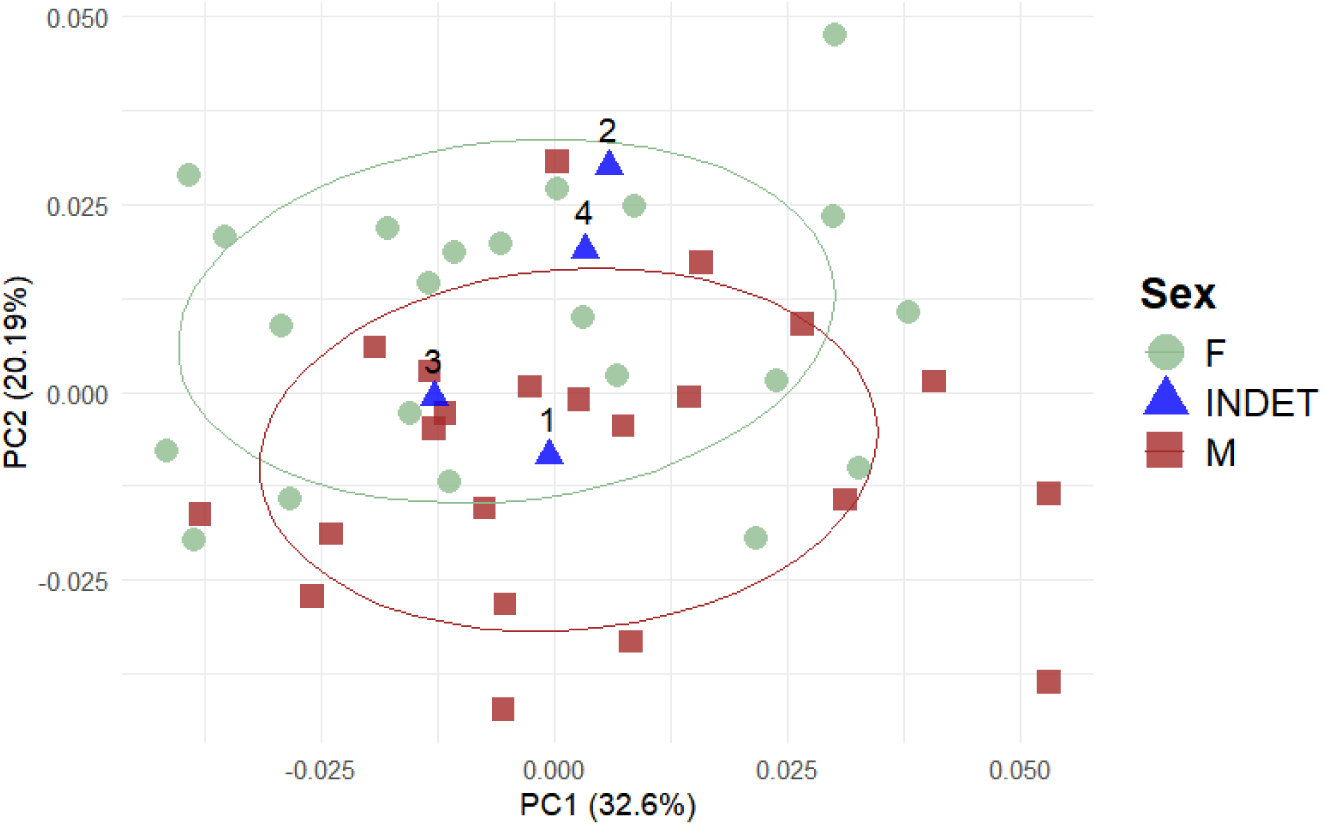
Morphospace is defined by the first two principal components from the PCA of the orbital shape. Contemporary male skulls are represented by brown squares, females by green dots, and ancient skulls of undetermined sex by blue dots. The ID number of each ancient subject is annotated next to the corresponding dot. A 68% confidence ellipse is associated with each of the male and female groups.

Within-group variability of our two ‘male’ and ‘female’ groups from the contemporary sample was assessed (Table 3). For each subject, the Procrustean distance between its shape and the mean shape of its group (mean male or female shape) is measured. Bartlett test was used to check whether male and female individuals had the same dispersion around their respective mean shape. Then a one-tailed Student’s t-test was performed to assess whether the intra-group variability of males was greater than the females one.

**Table 3.** Intragroup variability for males and females.

| Sex | Mean | Sd | Min | Max |
| --- | --- | --- | --- | --- |
| F | 0.0619 | 0.0108 | 0.0428 | 0.0786 |
| M | 0.0653 | 0.0148 | 0.0429 | 0.0934 |

Landmarks were grouped into 4 groups to form 4 regions (Fig. 7). The first one includes them all, representing the global facial shape. Then three specific facial regions were targeted: the orbits (zone 1 - n, mf_R, mf_L, fmo_R, fmo_L, e_R, e_L), the median axis including the piriform aperture (zone 2 - n, r, a, p, nl_R, nl_L), and the mid-face angulation (zone 3 - n, r, a, j_R, j_L).

**Fig 7.**
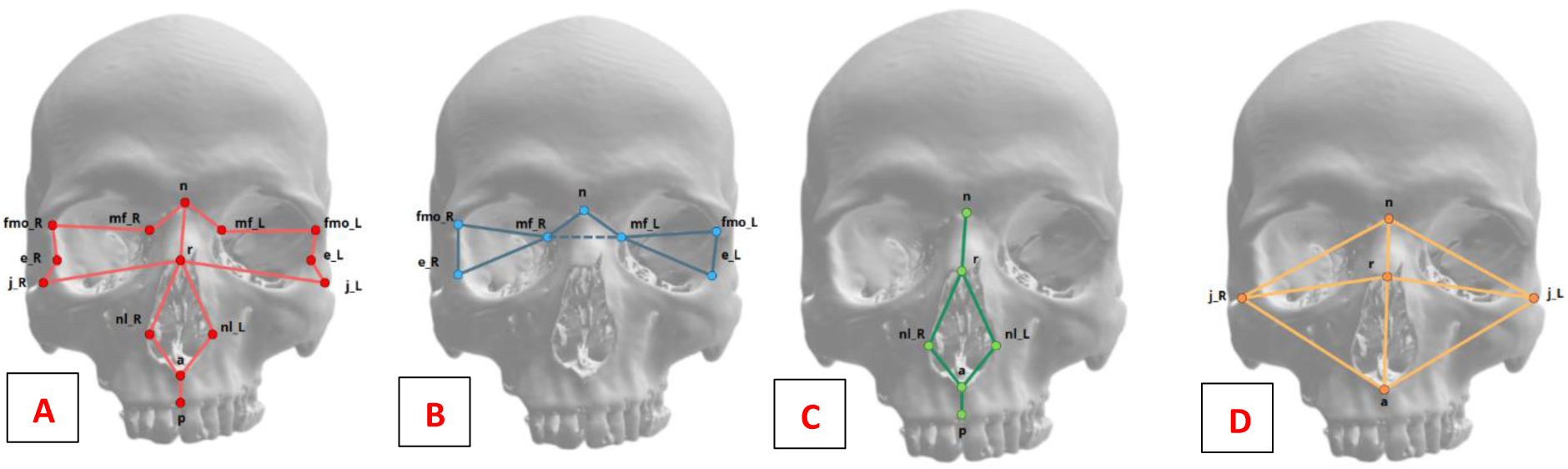
Anterior view of facial landmarks configuration and visualization of the shape associated with each zone drawn in two dimensions. (A) Zone ‘global’; (B) Zone 1 ‘orbits’; (C) Zone 2 ‘median axis’; (D) Zone 3 ‘mid-face angulation’

We applied a Thin-Plate Spline (TPS) transformation (Bookstein 1991), whose concept was first imagined by biologist D’arcy W. Thompson (Gould 1971), to visualize shape differences associated with facial sexual dimorphism. Thin-Plate Spline is an interpolation function that models the difference in shape between two landmark configurations by minimizing the bending energy required to deform a grid to a given landmark configuration (Cooke and Terhune, 2015). In the present study, it was performed on Procrustes data. Deformations were calculated in both two (x/y) dimensions and three (x/y/z) dimensions, as the 2D version facilitates visual comparison on flat projections, while the 3D approach faithfully reproduces volumes and avoids loss of information. First, the mean shape was calculated for each sex. The mean female shape was used as the reference and the mean male shape as the target for the transformation. The two average shapes were superimposed, projected onto a deformation grid and then the tps2d() (Fig. 8) or tps3d() (Appendix 1) functions from the R ‘Morpho’ package were applied. This procedure makes it possible to obtain the field of deformation vectors between the two shapes and to determine the amplitudes of deformation associated with each point on the grid. These amplitudes were then visualized using heatmaps, providing a synthetic and localized overview of the intensity of the deformations (Fig.9). The four predefined regions underwent the same process, allowing the deformation amplitudes specific to the facial areas to be identified and plotted.

**Fig 8.**
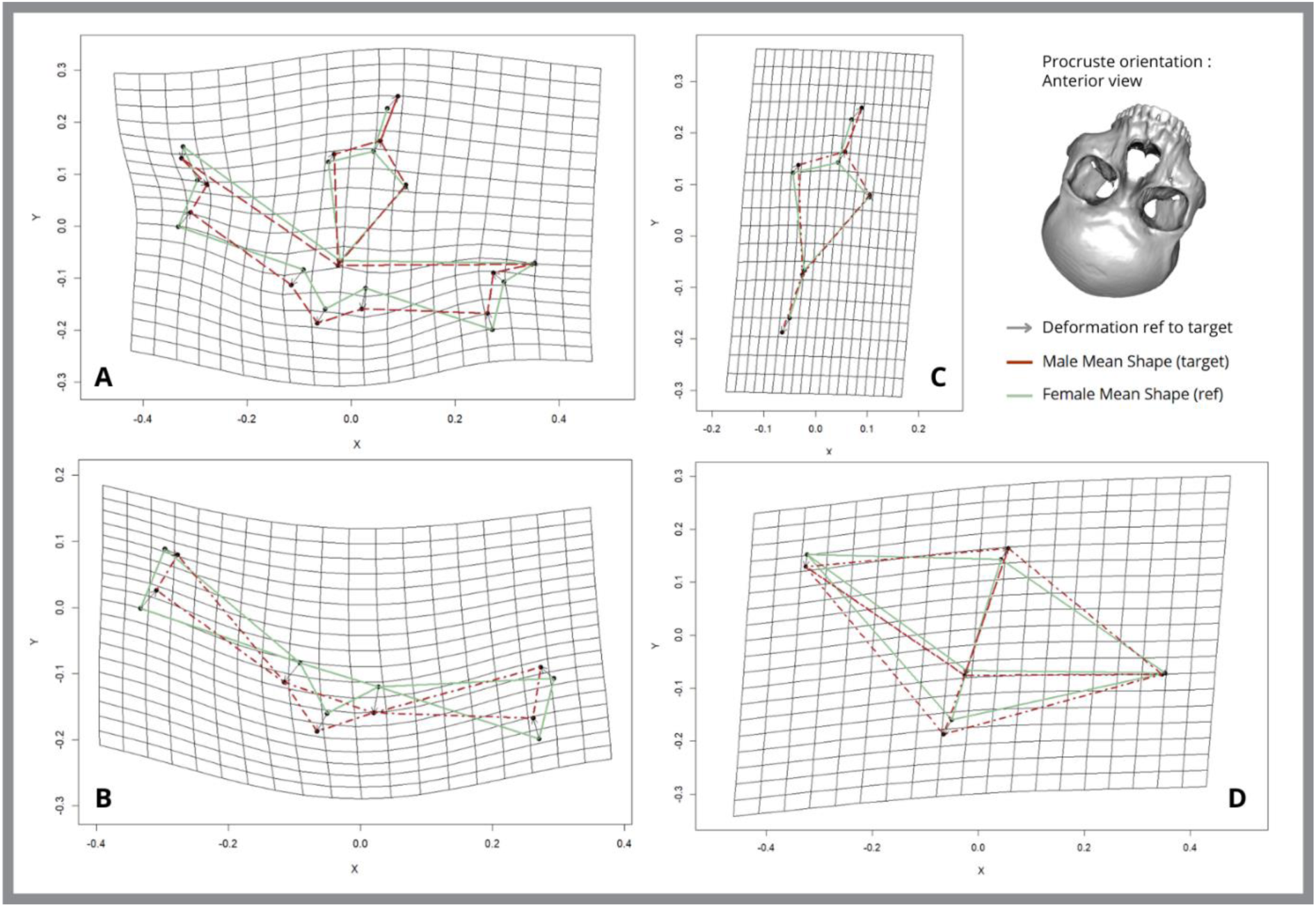
Thin-plate spline deformation grids in two dimensions depicting anterior view of mean female (green) and male (brown) facial shape. Those grids have been exaggerated three times to aid visualization. (A) Global Zone; (B) Orbits – zone1; (C) Median axis – zone2; (D) Mid-face angulation – zone3. Mean male shape (target) is represented in brown and mean female shape (reference) is plotted in green.

**Fig 9.**
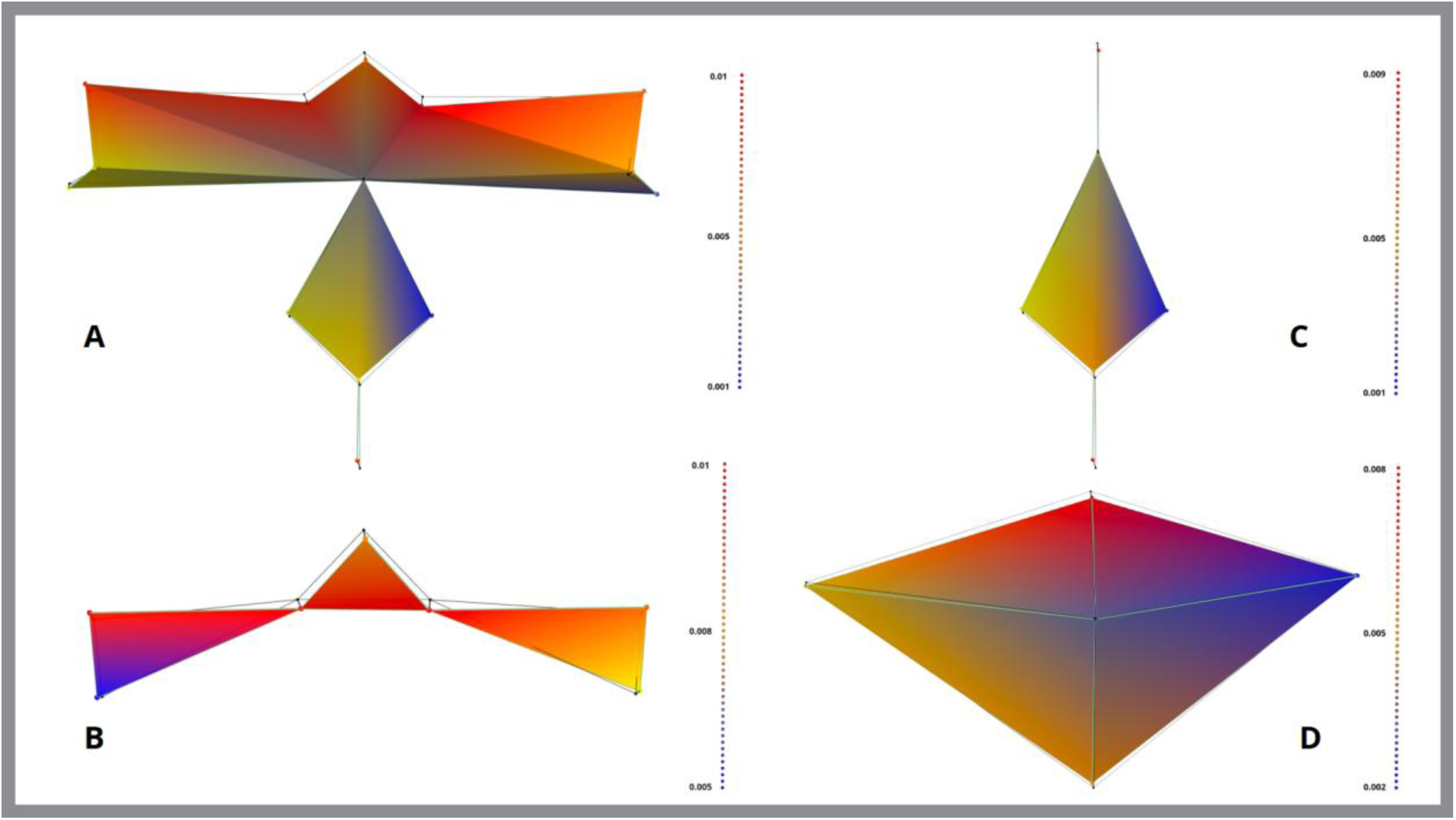
Anterior view of 3D heatmap showing deformation amplitudes between the female average shape (reference) and the male average shape (target). The values are represented by a gradient from blue to red: blue corresponds to low deformation amplitude values, and red to high deformation amplitude value s. (A) Global Zone; (B) Orbits – zone1; (C) Median axis – zone2; (D) Mid-face angulation – zone3.

To assess the significance of the shape differences between the groups of contemporary male and female subjects, Goodall’s F-test was performed using the R ‘geomorphs’ package (Table 4). This test compares the Procrustes distances, considering within -group variation. The analysis was first performed on the global shape and then on three predefined facial zones to account for potential differences between regions. A permutation procedure with 1000 iterations was used to compute the test.

**Table 4.** Goodall’s F-test performed on contemporary male and female groups for each facial region.

| Zone | R<sup>2</sup> | F | Z | p |
| --- | --- | --- | --- | --- |
| Global | 0,03914 | 1.71 | 1.60 | 0.048 * |
| Zone 1 | 0.0648 | 2.91 | 2.35 | 0.012 * |
| Zone 2 | 0.02718 | 1.17 | 0.59 | 0.274 |
| Zone 3 | 0.01916 | 0.82 | -0.05 | 0.515 |

Linear discriminant analysis (LDA) was performed to assess whether shape could efficiently predict sex. This analysis was carried out using the R ‘MASS’ package. Three models were created based on significant dimorphic facial regions. The first, called “Global”, is based on the fourteen facial landmarks. Given the ratio between the number of variables and the number of subjects, a prior dimension reduction by principal component analysis (PCA) was applied in order to respect the validity conditions - multivariate normality, homogeneity of the covariance matrices and multicollinearity. The first five principal components (PC1-PC5), which explained most of the variance, were retained. Likewise, the second model “Zone 1” is based on the first five principal components of a PCA computed

on the Procrustes coordinates corresponding to the orbital zone. The last model also concerns the Zone 1 but is directly based on the Procrustes coordinates (GPA) of the 7 landmarks defining the orbital region, without dimensional reduction, as it meets the validity conditions of the LDA (Table 5). To visualize the classification performance of the three discriminant models, ROC curves were computed for each configuration based on the Leave-One-Out Cross-Validation (LOOCV) results (Fig. 10), and the corresponding area under the curve (AUC) values were calculated to quantify the model performance.

**Fig 10.**
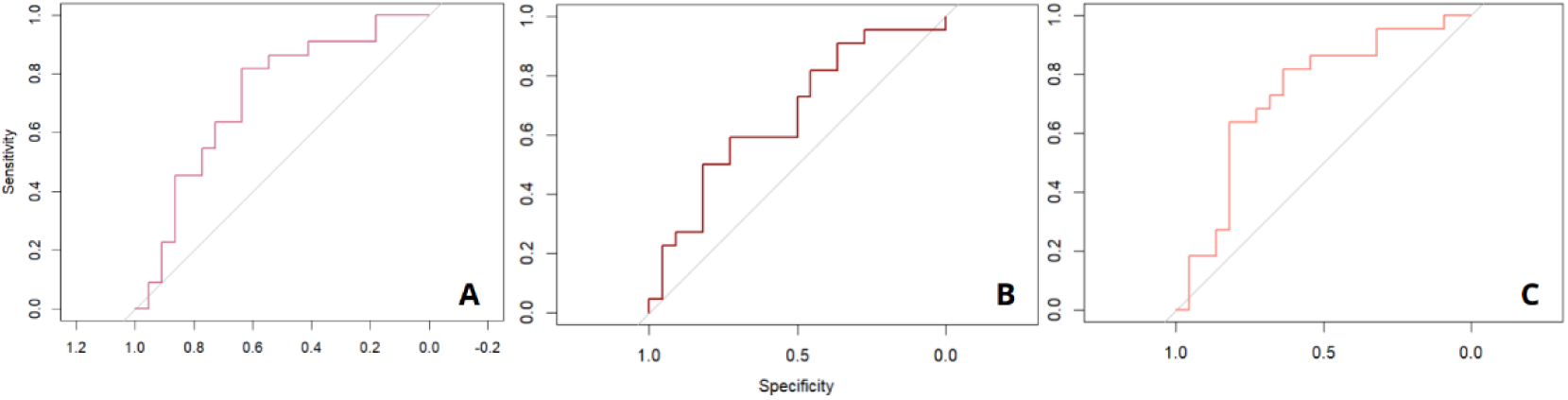
ROC curves illustrating the classification performance of the LDA models. (A) Global model based on PC1 –PC5 (AUC = 0.7190), (B) Zone 1 model based on PC1–PC5 (AUC = 0.6674), (C) Zone 1 model based on Procrustes coordinates (AUC = 0.7293). The diagonal grey line represents the random classification performance.

**Table 5.**
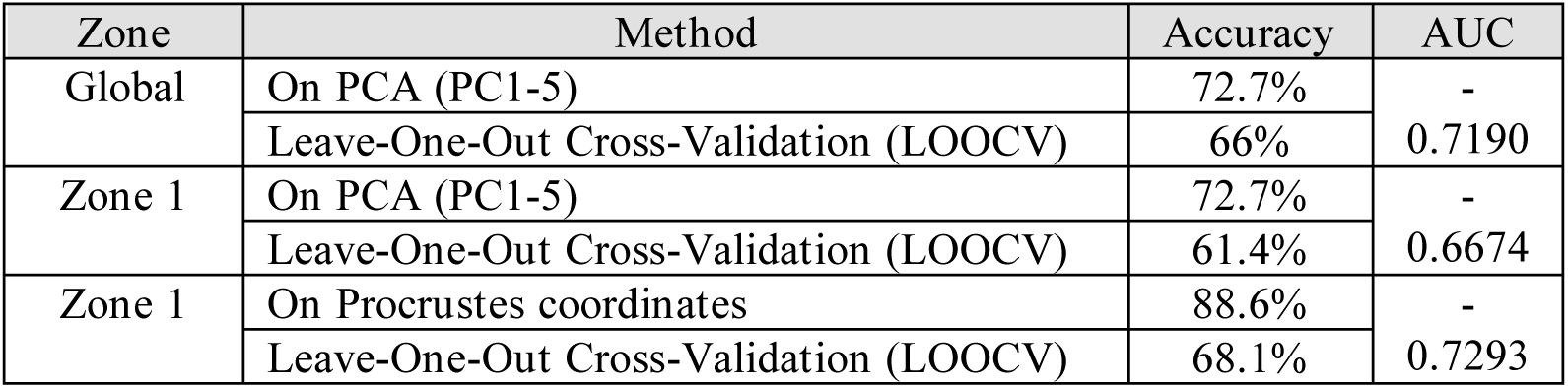
LDA models and sex classification accuracy.

### Sex prediction of the archeological sample

The Mahalanobis distances were calculated using the Mahalanobis distance function in R (Table 6). Based on the first five principal components retained from the principal component analysis of the orbital region (Zone 1) and the overall region (Global), the mean scores per landmark dimension and the covariance matrix were calculated separately for the male and female groups of the contemporary reference sample. For each sexually undetermined individual corresponding to the skulls found in the Grotte de La Médecine, the Mahalanobis distances were calculated with each group (male and female). These distances quantify the morphological proximity of each ancient individual to both groups of known sex (Slice & Ross, 2009). The closer the Mahalanobis distance is to one of the two groups, the more likely the individual’s shape is similar to this group. Thus, sex attribution was proceeded by assigning each ancient individual to the group (male or female) to which one showed the smallest Mahalanobis distance. However, these distances remain indicators of proximity without any proven significance or certainty in providing reliable sex estimation.

**Table 6.** Mahalanobis Distances.

| Zone | Subject | To male group | To female group | Proximity |
| --- | --- | --- | --- | --- |
| Global | 1 | 1.583475 | 3.499568 | M |
|  | 2 | 6.057865 | 3.363410 | F |
|  | 3 | 5.410390 | 5.375918 | F |
|  | 4 | 3.218660 | 1.109404 | F |
| Zone 1 | 1 | 1.163870 | 2.554274 | M |
|  | 2 | 12.632570 | 9.004608 | F |
|  | 3 | 7.916235 | 17.428489 | M |
|  | 4 | 5.839053 | 5.380280 | F |

To supplement this approach, the same discriminant models that had previously been applied to the contemporary sample were used on the archaeological sample. The predictions were based on PCA-reduced data for the overall facial region and Procrustes coordinates for the orbital region. Only the three-dimensional coordinates model was used for sex prediction, as it is the more reliable of the two models developed for Zone 1. This predictive approach enables a probabilistic classification (Table 7).

**Table 7.** Sex predictions.

| Zone | Subject | Probability M | Probability F | Prediction |
| --- | --- | --- | --- | --- |
| Global | 1 | 0.742 | 0.258 | M |
|  | 2 | 0.226 | 0.774 | F |
|  | 3 | 0.640 | 0.360 | INDET |
|  | 4 | 0.274 | 0.726 | F |
| Zone 1 | 1 | 0.960 | 0.040 | M |
|  | 2 | 0.023 | 0.977 | F |
|  | 3 | 0.660 | 0.340 | M |
|  | 4 | 0.440 | 0.560 | INDET |

## Results

### Landmarks repeatability

For the intra-observer variability, mean Euclidean distances ranged from 0.70 mm (r) to 1.47 mm (e_L), and mean error percentage ranged from 0.20% to 0.41% whereas for the inter-observer variability, mean Euclidean distances ranged from 0.79mm (a) to 3.59mm (n), and mean error percentages ranged from 0.20% to 1.07%. Regarding landmark placement repeatability, intraclass correlation coefficients (ICC) were consistently high across all landmarks. The lowest ICCs were observed for the maxillofrontal landmarks (intra-observer) and the nasion (inter-observer), while the highest ICCs were found for the prosthion landmarks (intra-observer) and the rhinion landmark (inter-observer) (Table 2).

### Preliminary analysis

The principal component analysis showed that the first two components (PC1 and PC2) describe 42,92 % of the total morphological variability of the face considering the shape (PC1 = 24,5 %, PC2 = 18,42 %). There is a large overlap between female (F) and male (M) individuals along the PC1 and PC2 (Fig. 5).

The test for homogeneity of variances revealed no significant difference in shape dispersion between males and females (Bartlett’s K-squared = 1.9806, df = 1, p-value = 0.1593). This indicates that both groups have comparable levels of within-group morphological variability around their respective mean shapes. The one-tailed Student’s t-test did not reveal significantly greater intra-group variability in males compared to females (t = 0.86921, df = 42, p-value = 0.1948).

### Zone-specific shape dimorphism

Deformation amplitudes over the entire facial form (Global) vary between 0.001 and 0.01, with higher values observed in the nasion, frontomallar orbital, maxillofrontal and prosthion, where the amplitudes range from 0.008 to 0.01. For the orbital zone (Zone 1), deformation amplitudes range from 0.005 to 0.01. Most of its landmarks show deformation values greater than 0.007, except for the ectoconchions, which show lower amplitudes. For the medial axis (zone 2), the deformation amplitudes vary between 0.001 and 0.009. Only the nasion and prosthion keep amplitudes values greater than 0.008. Finally, in the mid-face angulation shape (Zone 3), the amplitudes range from 0.002 to 0.008. The rhinion, acanthion and jugal landmarks have values below 0.005. Zones 2 and 3 appear to contain the least deformed landmarks, whereas zone 1 contains the most deformed ones (Fig. 8 & 9). Visual analysis of the three-dimensional superimpositions of the male and female mean shapes reveals differences in the relative landmarks configuration. The maxillofrontals and nasion are in a ‘superior’ position in the male average shape compared to the female. The ectoconchions, nasolaterals and frontomallar orbitals appear to be displaced medio-distally in the male shape. The rhinion seems to be quite similar between the two configurations, although the male shape shows a slight anterosuperior translation compared to the female mean shape. Finally, the prosthion and acanthion are more ‘distal’ in the male than in the female average shape.

The results of Goodall’s F-test showed only 2 significant zones (Table 4). For the “Global” zone, the analysis indicates a significant difference between the male and female facial shapes (F = 1.71, Z = 1.60, p = 0.048), with a coefficient of determination R² = 0.03914, which means that almost 4% of the total shape variance is explained by sex. The division into regions makes it possible to specify which part of the face is most involved in facial sexual dimorphism. Zone 1 (orbital region) shows a significant sexual dimorphism and explains almost 6.5% of the total shape variance (F = 2.91, Z = 2.35, p = 0.012, R² = 0.0648). On the other hand, neither Zone 2 (median axis; p = 0.274) nor Zone 3 (mid-face angulation; p = 0.515) showed significant shape dimorphism.

Only regions exhibiting significant sexual dimorphism were included in the subsequent analyses, therefore excluding zone 2 and zone 3.

The principal component analysis of the orbital zone showed that the first two components (PC1 and PC2) describe 52,79 % of the total morphological variability of the orbits considering the shape (PC1 = 36.2 %, PC2 =20,19 %) (Fig. 6). The PCA scores revealed that the second principal component (PC2) shows a significant sex-related difference between male and female groups (W = 365, p = 0.004). The analysis of variable contributions to PC2 indicates that the nasion and the maxillofrontals landmarks contribute most prominently to the morphological variation captured along this axis within the orbital region. Chalcolithic subjects 2 and 4 appear to align more closely with the female distribution, whereas subjects 1 and 3 are positioned closer to the male distribution. Nevertheless, subject 3 occupies a more intermediate position compared to subject 1.

### Sex classification

The results from linear discriminant analysis (LDA) performed on three models based on significant dimorphic facial regions showed that the ‘Global’ and the ‘Zone 1’ models both based on PCA scores achieved a classification accuracy of 72.7% (CI 95 % = [0.5721, 0.8504]). While the “Zone 1” model based on three-dimensional coordinates corresponding to the orbits reaches 88.6% (CI 95 % = [0.7544, 0.9621]). In the latter, 86.4% of females and 90.9% of males from the contemporary population were correctly classified. In leave-one-out cross-validation (LOOCV), the accuracy of Zone 1 model, based on Procrustes coordinates drops to 68.1% while in the Global model, it decreases to 66%. The third model, “Zone 1” based on the first five principal components (PC1-PC5) drops to 61.4% using leave-one-out cross-validation (LOOCV). The highest discriminative ability was observed in the “Zone 1” model based on three-dimensional coordinates (AUC =0.7293), followed by the “Global” model (AUC = 0.7190) and finally the “Zone 1” model based on PCA scores (AUC = 0.6674) (Table 5; Fig.10).

### Sex prediction of the archeological sample

The Mahalanobis distances show, firstly for the overall facial shape (Global) that the subject 1 was closer to the male average shape (d² males = 1.58) than to the female average shape. In contrast, 2, 3, and 4 were closer to the female mean shape, especially subject 4 (d² females = 1.11; d² males = 3.22). Secondly, in the orbital region (Zone 1), a similar pattern was observed for subject 1, 2, and 4: 1 remained closer to the male mean shape, while subjects 2 and 4 were closer to the female mean shape. Subject 3 however showed a shift, being closer to the female average shape in the overall shape but closer to the male average shape in Zone 1 (d² males = 7.92; d² females = 17.43) (Table 6).

The classification threshold for applying sex prediction models to ancient individuals has been set at 65%. Below this value, the probability of belonging to one sex or the other isn’t considered valuable, which classifies the sex of the individual as undetermined. In the overall model (Global), individuals 1 and 2 are classified as male (74.2%) and female (77.4%) respectively. Individual 3 is not assigned to any sex because its probabilities are below the required threshold. Individual 4 is classified as female (72.6%). In the orbital region (Zone 1), individuals 1 and 2 are classified as male (96%) and female (97.7%) respectively, as it was in the first model. Individual 3 was barely above the threshold (66%) and is therefore classified as male. The sex of individual 4 remains undetermined in this model (Table 7).

## Discussion

Virtual anthropology enables the study of three-dimensional morphological structures using data represented in a computer environment (Weber et al. 2001). These datasets can be acquired using a variety of methods. The most common ones are computed tomography (CT) scanning, surface scanning, and photogrammetry (Waltenberger et al. 2021). These methods can be employed on both ancient and modern samples, as illustrated by the use of a post-mortem CT in the present study. Using imagery enables access to hidden structures, to generate 3D models that will remain usable and shareable with the rest of the scientific community, and to study volumes, areas, curves, or points data to obtain maximum information (Weber 2014). Virtual anthropology is particularly relevant for the study of anatomically inaccessible structures, where imaging is essential to access and study skeletal remains, such as the *Grotte de La Médecine* case. This geometric morphometrics study focused on shape was based on 3D landmarks coordinates placed on the face of human skulls. Analysis performed on virtual objects, particularly those based on landmark settings, have been shown to be reliable and valid for enriching morphoscopic and morphometric studies (Richtsmeier et al. 1995; Hale et al. 2014). Nevertheless, differences in landmark placement between two or more observers usually occur, as demonstrated in several studies (Shearer et al. 2017; Robinson 2017; Fox et al. 2020), which showed that inter-observer error was generally higher than errors among an observer. Moreover, unlike intra-observer error, inter-observer error doesn’t necessarily decrease with increased experience in geometric morphometrics. Most importantly these studies showed that the inter-observer error could bias the results of statistical shape variation analyses. Using type I landmarks (Bookstein 1990) doesn’t guarantee better repeatability (Wärmländer et al. 2018). Their research shows that the landmarks typologies I, II or III don’t influence their repeatability, and therefore the robustness of the associated results, more than the types of measurement and the measurement tools used. Depending on the research protocol, type I (biological landmarks) isn’t necessarily more reliable than constructed landmarks. Nevertheless, in our study, type I landmarks were used. According to the Intraclass Correlation Coefficient, there is a high intra- and inter-observer agreement in the coordinates of the landmarks used to describe craniofacial shapes in this study.

The first objective of this research was to apply geometric morphometric methods to quantitatively describe facial sexual dimorphism within a contemporary French population. Based on a sample of 44 adult skulls of known sex, the principal component analysis (PCA) performed on the full set of facial landmarks revealed substantial overlap between male and female individuals. In contrast, the orbital region (zone 1) provided a clearer distinction between the sexes. A Student’s t -test conducted on this PCA (zone 1) indicated a significant difference in PC scores between males and females on PC2. This suggests a specific role of the orbital region in facial sexual dimorphism. Upon examining the eigenvector coefficients, the landmarks contributing most to the second principal component are the maxillofrontals (right and left) and the nasion. According to the results of the Thin-plate spline transformation based on the male (target) and the female (reference) average shapes, the greatest deformation amplitudes are indeed associated to the landmarks defining the orbital region: nasion, frontomalare orbital (right and left), maxillofrontal (right and left), and ectoconchion (right and left). These observations were supported by Goodall’s F-tests, which revealed significant shape differences between male and female individuals, both at the global scale and within the facial regions analyzed separately. The orbital region, as expected from exploratory analysis, stands out as an anatomically dimorphic area. In contrast, the relatively weak, yet significant global zone signal, may be explained by the diluting effect of non-dimorphic structure. The median axis (zone 2) and mid-facial angulation (zone 3) were not found to be significantly dimorphic between sexes, which may attenuate the influence of sexual dimorphism on overall shape variation when all facial regions are jointly analyzed. The absence of statistically significant results in zone 2 diverge from morphometrics findings on Brazilian and Indian samples that show the nasal bone and piriform aperture were significantly different between male and female (Cantin et al. 2009; Singh 2020). This difference had also been highlighted on the piriform aperture and subnasal region by Thin Plate Spline analyses conducted on a Portuguese sample (Rosas & Bastir 2002).

The second objective was to develop and evaluate discriminant models based on facial regions exhibiting significant sexual dimorphism in the contemporary French sample. The highest accuracy of 88.6% was based on the Procrustes coordinates of the orbital region. Accuracy was found to be 86.4% for females and 90.9% for males. Following a leave-one-out cross-validation (LOOCV) procedure, this value decreased to 68.1%. The classification performance (AUC) of the orbital region was superior to the global facial shape defined by all 14 landmarks, for which the model yielded a classification accuracy of 72.7%, 68.18% for females and 77.27% for males with a LOOCV accuracy around 66%. We compared our results with previous research identifying the orbital region as significantly discriminant for sex classification across different population samples. In a Portuguese sample of 125 adult individuals who died in the 20th century, sex estimation based on geometric morphometric analysis of orbital shape yielded an accuracy of 64.15% for females and 60.75% for males (Gonzalez et al. 2011). Another study conducted on 211 adult skulls from a Bosnia concluded, through discriminant function analysis (DFA), that sex classification based solely on the shape of the orbital region achieved an accuracy of 82.01% for males and 80.55% for females (Ajanović et al. 2023). Regarding research conducted on 139 skulls from Central European subjects, classification models based on three-dimensional coordinates reached an accuracy of 74% (Bigoni et al. 2010). These results are consistent with descriptions of sexual dimorphism in the orbital region, which characterize female orbits as more oval-shaped, whereas male orbits tend to be more angular (Pretorius et al. 2006). As for discriminant analyses based on measurements, a study examined 176 skulls from a Greek sample. It reported a classification accuracy of 79% based on the width of the orbital region (Chovalopoulou et al. 2016). Likewise, the accuracy based on orbital shape was found to be 78.7% for females and 70.8% for males in an East Indian sample of 118 skulls studied (Sarkar & Mukhopadhyay 2018). In another study based on 118 american subjects of both white and black, the classification accuracy was 73.3% for females and 80.0% for males among white subjects, and 79.31% for females and 75.86% for males among black subjects (Kimmerle et al. 2008). Their findings appear to align with earlier studies conducted on an other american sample of 247 skulls which employed discriminant functions based on cranial measurements (Giles & Elliot 1963). These studies support the idea that the orbital region provides anatomical information that can be used to estimate sex across diverse populations. Although accuracy rates vary depending on sample size, origin, and methodological approaches. Ranging from traditional craniometric measurements to geometric morphometric methods, the orbital region consistently emerges as a particularly reliable contributor to sexual dimorphism. The results obtained from our French sample further confirm the usefulness of the orbital region in sex prediction models employed for the biological profile assessment in forensic and biological anthropology.

Finally, this study aimed to assess whether the regions exhibiting significant sexual dimorphism could be used to predict the sex of ancient individuals, using discriminant models developed from the facial shapes of the contemporary French sample. The projection of chalcolithic subjects into the principal component spaces defined by the reference sample shows different distributions. In the global morphometric space, the strong overlap between groups limits interpretation, as the subjects don’t visually cluster close to either sex. However, in the PCA based on the orbital region, subject 1 is positioned closer to the male distribution along PC2, while 2 and 4 are nearer to the female distribution. The third subject is positioned between the two groups. The projection enables a preliminary comparative assessment of their facial morphology. Their distribution in morphospace suggests a heterogeneity of shapes between them, with some individuals showing affinities with contemporary distribution. The use of Mahalanobis distances (D²), calculated relative to the contemporary male and female mean shapes, provide a quantitative measure of morphological proximity. These distances were further supported by a probabilistic approach using LDA models, which estimated the likelihood probabilities of each ancient individual belonging to either sex. The subject 3 appears morphologically intermediate, displaying comparable probabilities of classification into both groups. Subject 1 was associated with a high probability of belonging to the male sex. In contrast, 2 and 4 were associated with the female sex. Subject 2 showed high probabilities, while 4, despite an acceptable probability in the global model, was not classified by the orbital region model. It is important to acknowledge that this approach is limited by morphological changes that have occurred since the Neolithic period. A study conducted on a male Chinese sample, including 161 Neolithic, 423 from the Bronze Age, and 134 modern subjects, examined the evolution of craniofacial morphology from the Neolithic period to the present day. Based on 21 cranial measurements, the authors identified 14 traits that showed statistically significant changes over time. Nine traits decreased significantly, while three increased. The most marked variations concerned the nasal index (−8.2%), the orbital index (+6.9%), and nasal breadth (−6.0%) (Wu et al. 2007). Although morphological changes vary across populations (Brown et al. 2004), as does the expression of sexual dimorphism (Garvin 2012; Ubelaker & DeGaglia 2017; Ubelaker & DeGaglia 2020). This approach presents promising prospects. It could ultimately serve as a basis for evaluating the reliability of sex predictions from discriminant analyses by comparing results across ancient and contemporary samples, provided that new chalcolithic human remains are discovered in France. Building on this approach, future research could further investigate the influence of age on facial morphology. Numerous studies have documented age-related morphological variations, particularly in the orbital region, the mandible, and the cranial vault (Behrents 1985; Enlow & Hans 1996; Doual et al.,1997; Patterson et al. 2007; Avelar 2017). Such variations may constitute a substantial source of shape variability and act as confounding factors that compromise the performance of discriminant models, especially the bone resorption with aging (Walsh 2015). Accounting for age as a variable could therefore enhance the study of contemporary morphological shapes and improve the accuracy of predictive models when applied to ancient subjects.

## Conclusion

This study highlights the relevance of virtual anthropology and geometric morphometrics in the analysis of facial sexual dimorphism. Based on a contemporary French sample, the results reveal a significant shape difference between males and females. The orbital region emerges as a discriminant area within this sample. These findings are consistent with those observed in other populations, confirming the orbital region as a relevant anatomical area for sex estimation. Although preliminary, the application of these models to chalcolithic subjects underscores the need for further research to refine our understanding of the evolutionary dynamics of craniofacial morphology over time. Moreover, this approach raises methodological considerations regarding the validity and limitations of using contemporary reference datasets in bioarchaeological interpretations. Overall, this study contributes to a better understanding of human sexual dimorphism and aligns with ongoing efforts to improve sex estimation methods in both forensic and archaeological contexts. It emphasizes the value of 3D imaging and landmark-based geometric morphometrics for shape analysis in biological anthropology.

## Acknowledgments

We are grateful to the University of Toulouse for supporting this research, and to the Musée de Millau et des Grands Causses (MUMIG) for granting access to the skeletal remains from the Grotte de La Médecine.

## Data availability

Data supporting the results of this study can be obtained from the corresponding author upon request.

